# Bees Without Flowers: Before Peak Bloom, Diverse Native Bees Visit Insect-produced Honeydew Sugars

**DOI:** 10.1101/082271

**Authors:** Joan M. Meiners, Terry L. Griswold, David J. Harris, S.K. Morgan Ernest

**Affiliations:** School of Natural Resources and Environment, University of Florida, Gainesville, FL 32611.; USDA-ARS Pollinating Insects Research Unit, Utah State University, Logan, UT 84321.; Department of Wildlife Ecology and Conservation, University of Florida, Gainesville, FL 32611.

**Keywords:** native bees, non-floral foraging, honeydew sugars, foraging behaviors, social cues, bee communities

## Abstract

Bee foragers respond to complex visual, olfactory, and extrasensory cues to optimize searches for floral rewards. Their abilities to detect and distinguish floral colors, shapes, volatiles, and ultraviolet signals, and even gauge nectar availability from changes in floral humidity or electric fields are well studied. Bee foraging behaviors in the absence of floral cues, however, are rarely considered. We observed forty-four species of wild bees visiting inconspicuous, non-flowering shrubs during early spring in a protected, Mediterranean habitat. We determined experimentally that these bees were accessing sugary honeydew secretions from scale insects without the aid of standard cues. While honeydew use is known among some social Hymenoptera, its use across a diverse community of mostly solitary bees is a novel observation. The widespread ability of native bees to locate and use unadvertised, non-floral sugars suggests unappreciated sensory mechanisms and/or the existence of a social foraging network among solitary bees that may influence how native bee communities cope with increasing environmental change.

## Introduction

Bees and flowers are inextricably linked. Their mutualistic relationship has been a timeless focus for poets, artists and naturalists, as well as field ecologists, behavioral scientists, and evolutionary biologists. The obsession is not without merit. Bee visits to flowers for nectar and pollen are so crucial to angiosperm reproduction that bee preferences for floral colors, shapes, and scents have been credited with driving floral trait evolution, a radiation in angiosperm species diversity during the Late Cretaceous, and current plant community composition (Regal 1977; Ohashi and Yahara 2001; Wright and Schiestl 2009; de Jager et al. 2011; Ollerton et al. 2011; Bukovac et al. 2016). Because of this influential mutualism, research on bee foraging has focused on how bees detect and respond to floral visual and olfactory cues, petal thermal signatures, humidity signals from nectar reserves, and even floral electric fields (Herrera 1995; Chittka et al. 1999; Dyer et al. 2006; Whitney et al. 2008; Wright and Schiestl 2009; de Jager et al. 2011; Clarke et al. 2013; Frisch 2014; Orbán and Plowright 2014). Very little, however, is known about bee foraging behaviors in the absence of floral cues, particularly among wild, solitary bee species.

Bees require sugar, usually as floral nectar, and protein, typically from pollen, for energy and reproduction (Michener 2007). While specialist bee species are particular about their pollen sources, bee visits to flowers for nectar sugars are usually indiscriminate (Linsley 1958).

Honeydew is a nectar-like carbohydrate-rich excretion produced as a feeding by-product by phytophagous Hemipterans, such as scale insects (Hemiptera: Coccoidea) and aphids (Hemiptera: Aphididae), that can sometimes be more nutrient-rich than floral nectar (Batra 1993). Some insects, most notably ants, attain increased fitness and longevity by using honeydew as an additional sugar source (Zoebelein 1957; Wäckers et al. 2008; Koch et al. 2011).

Honeydew use among bees, while digestively plausible and potentially broadly advantageous given global concern about bee declines and their temporal isolation from host flowers (Potts et al. 2010; Bartomeus et al. 2011), has been only sparsely documented, usually as isolated occurrences, and almost exclusively among social, colonial species (Santas 1983; Crane and Walker 1985; Batra 1993; Bishop 1994; Konrad et al. 2009; Koch et al. 2011).

Widespread use of honeydew by diverse solitary bee species would have interesting implications for bee ecology, behavior, and conservation for two important reasons: 1) it represents a departure from the classic paradigm of the bee-flower mutualism as a tightly coupled relationship, and 2) it suggests an as-yet unstudied source of resilience, behavioral and physiological, among bees foraging to survive in a changing climate. Honeydew as a sugar compound is non-volatile, colorless, does not fluoresce or absorb UV, and occurs independently of flowering resources (Thorp et al. 1975; Friel et al. 2000; Frisch 2014). Prior to blooming of the host plant, therefore, it is a resource without apparent visual, olfactory, or floral advertisement. An ability of bees to expand conventional search images and diet breadth to include resources such as honeydew could be an important adaptation in habitats, like Mediterranean biomes, where the flora is predicted to be especially sensitive to global change (Klausmeyer and Shaw 2009). Faced with increasingly unpredictable foraging scenarios, honeydew could be an invaluable emergency resource for bees who are able to find it.

Working in the Mediterranean, chaparral habitats of Pinnacles National Park in the Inner South Coast Range of California, one of us (J. Meiners) observed a diverse array of native, mostly solitary bee species visiting large, woody, pre-bloom *Adenostoma fasciculatum* shrubs (Rosaceae) during the early spring when floral resources were still very limited (Fig. 1). Some of these shrubs were covered in a dark ‘sooty mold,’ known to grow on the honeydew excretions of scale insects (Hemiptera:Coccoidea) (Santas 1983; Crane and Walker 1985; Wäckers et al. 2008). To evaluate this perplexing attraction to moldy plants, we began noting the mold and bloom condition of each *A. fasciculatum* shrub every time we collected a bee from these plants during sampling for a broader biodiversity survey. Surprisingly, we recorded nearly four times as many bees visiting moldy, pre-bloom *A. fasciculatum* individuals as visited mold-free varieties, or either mold condition after flowering, confirming the association of bees with mold but raising new questions about the appeal and mechanism (Fig. 2). These results prompted us to design an experiment for the following early spring to evaluate three central questions: 1) Why are bees visiting these pre-bloom plants?; 2) What are the potential visual, olfactory, thermal, or insect-insect cues alerting bees to this resource?; and 3) How widespread is this behavior across the bee community?

**Figure 1:**
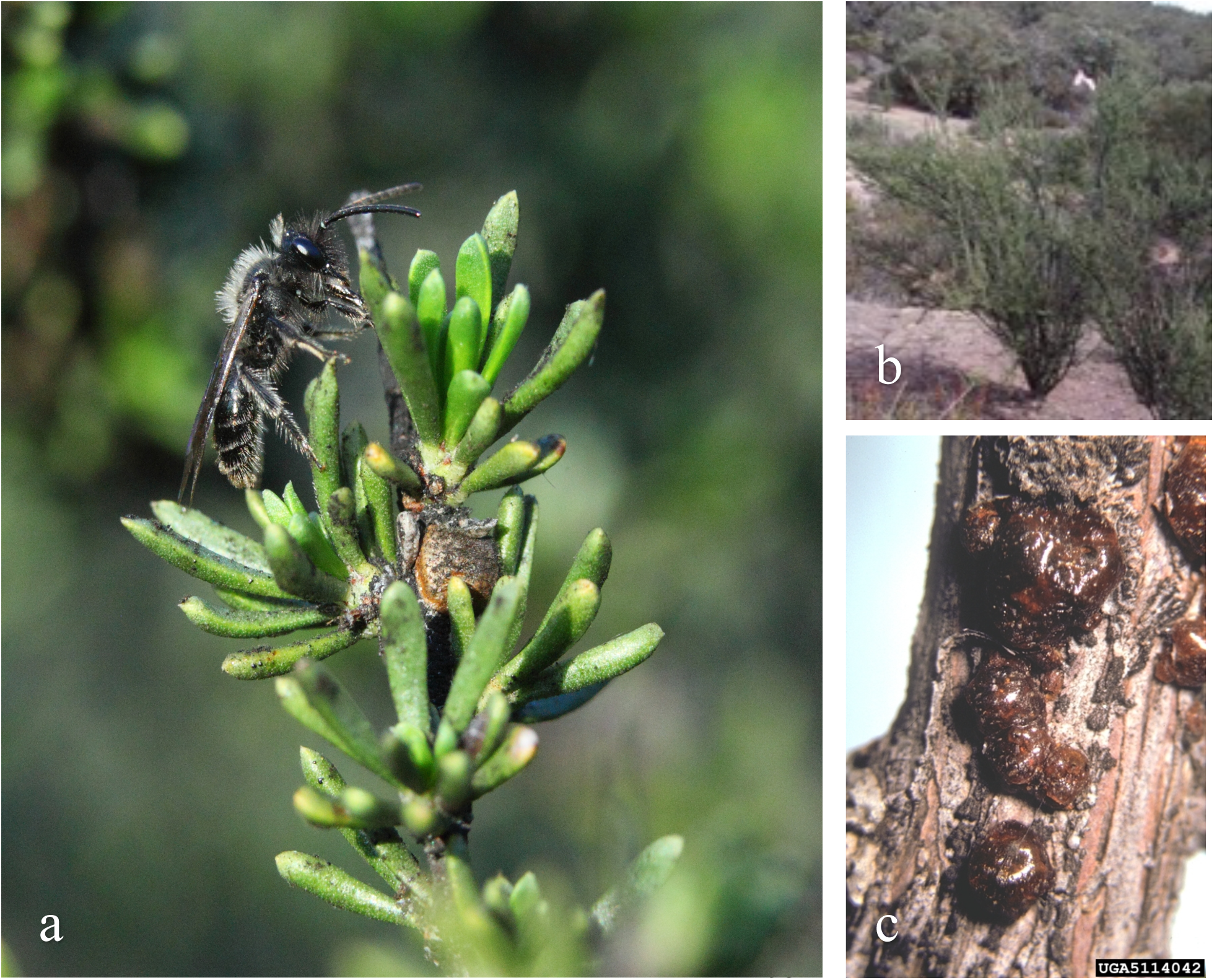
Images of organisms involved in the described system. (a) Native bee (*Andrena* sp.) foraging on a moldy, non-flowering *Adenostoma fasciculatum* shrub (left, Photo Credit Paul Johnson, NPS), (b) a typical *A. fasciculatum* shrub in a pre-bloom Pinnacles landscape (top right, J. Meiners), and (c) an image of lac scales (*Tachardiella* sp.) on an *A. fasciculatum* branch (bottom right, Creative Commons United States National Collection of Scale Insects Photographs, USDA Agricultural Research Service, Bugwood.org).

**Figure 2:**
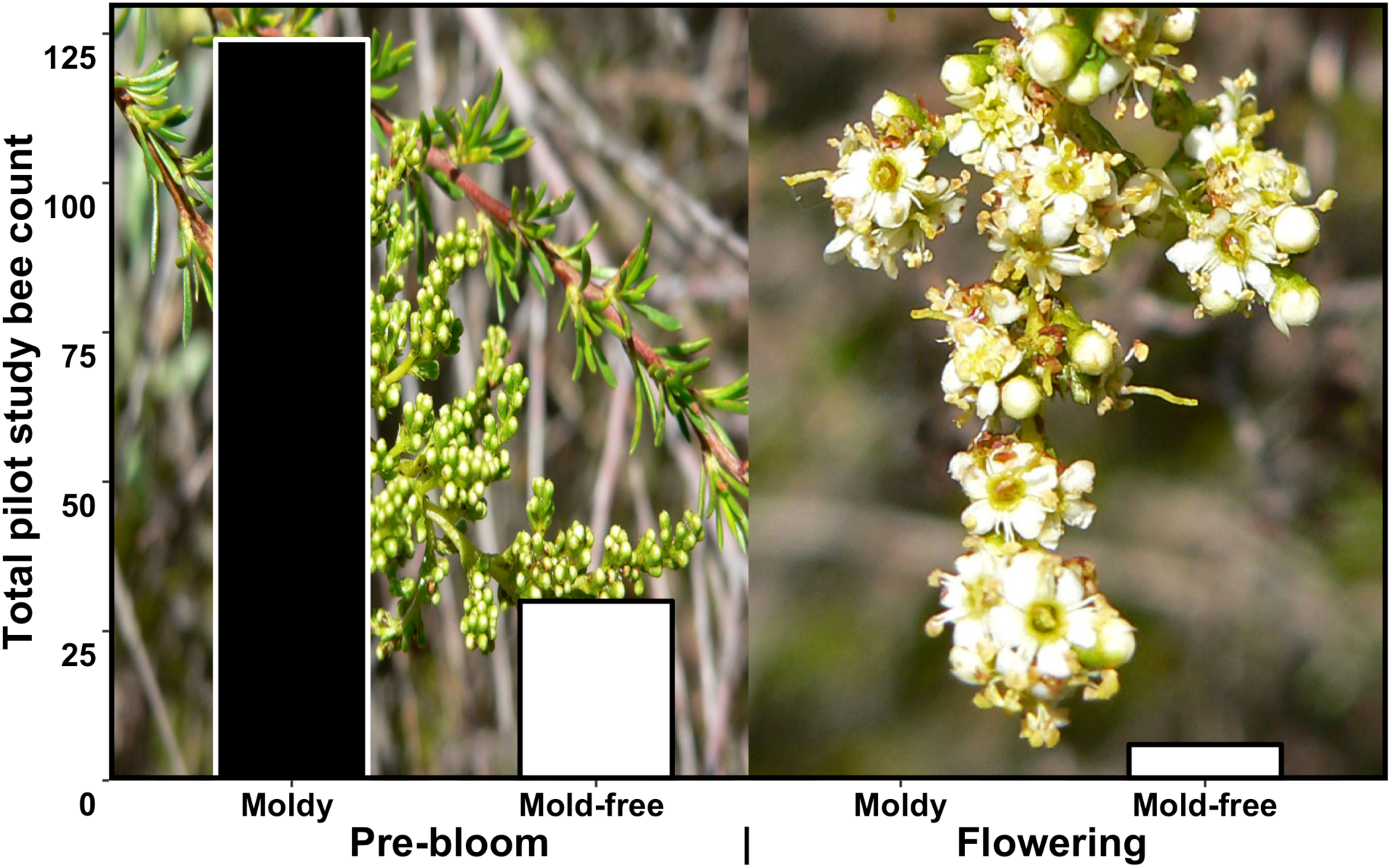
Number of bees collected on different *Adenostoma fasciculatum* plant conditions: pre-bloom (left panel), flowering (right panel), moldy (black bars) and mold-free (white bars). Between February and June during the pilot year of study, we collected a total of 160 native bees visiting *A. fasciculatum* shrubs, 124 of which were on non-flowering, moldy plants, and 30 of which were visiting plants without mold or flowers. After flowering commenced, we did not collect a single bee visiting plants with mold and flowers, and only six bees on non-moldy shrubs in bloom, perhaps because other floral resources are preferred at that time. Based on these results, we conducted our study during the period prior to *A. fasciculatum* flowering. Photos by Stan Shebs, CC BY-SA 3.0, https://commons.wikimedia.org/w/index.php?curid=2407692.

## Materials and Methods

### Experimental Design

We designed seven experimental treatments to differentiate the possible mechanisms and causes of bee attraction to sooty mold, and randomly assigned them to 'naturally moldy' and 'mold-free' *Adenostoma fasciculatum* shrubs at three distinct 1-hectare experimental sites in natural areas within Pinnacles National Park in San Benito County, California. Each selected site was dominated by the large, hardy, allelopathic *A. fasciculatum* shrubs and included a mixture of shrubs of similar stature that we could designate as either 'mold-free' (absent of sooty mold and scale insects) or 'naturally moldy' (visibly covered on more than 50% of branches by sooty mold). We applied each treatment (outlined below and in Table A1) to three woody shrubs of pre-bloom *A. fasciculatum* at each of the three sites, for a total of nine shrub replicates for each of seven treatments.

To control for any attraction, reflectance, or humidity signal of moisture, all seven treatments consisted of 5 ounces of fluid sprayed on the assigned shrub, as follows:

Naturally-moldy plants were sprayed with either (i) water to assess baseline bee visitation to moldy plants, or (ii) a natural, short-residual insecticide (*Orange Guard*^®^ Water Based Indoor/Outdoor Home Pest Control, active ingredient d-Limonene, 5.8%) to evaluate the influence of live scale insects on bee visitation by halting their activity, while leaving sugars and mold intact.

Mold-free plants were sprayed with either (i) water to quantify random bee visitation, (ii) insecticide to test for an effect of this chemical on bee activity, (iii) non-toxic black paint to test for an attraction to either the dark visual cue of mold or to potentially higher branch surface temperatures, which recent research has found to be attractive to bees (Dyer et al. 2006), (iv) a colorless, odorless 20% 1:1 Sucrose:Fructose solution mixed from chemical-grade sugars to mimic the composition of insect honeydew (Wäckers et al. 2008), or (v) a combination of both the black paint and the sugar mixture to simulate the attraction of natural mold and examine interaction effects (treatments summarized in Table A1).

### Sampling Protocol

Because the pilot study indicated that bee visits to honeydew were restricted to the early season (Fig. 2), we concentrated our experiment in the period before peak bloom. We visited each site three times between late February and late April, when native bee activity has begun at Pinnacles National Park but prior to peak bloom of the plant community. Sampling was conducted at one of the three sites per week on calm, sunny days over 15°C to ensure adequate bee activity. At 9am on each sampling day, we began by refreshing all plants with 5oz of their assigned treatment spray, which remained the same throughout the experiment. After waiting an hour for the effect of the short-residual insecticide to take place and subside, and for bee activity to approach peak levels for the day, a randomly ordered shrub list was divided between two collectors, who spent five minutes sequentially netting all bees visiting each respective plant. Temperature, wind speed, humidity, barometric pressure, and an estimate of cloud cover were recorded every thirty minutes during sampling. We sampled all twenty-one plants at a site once in the morning, beginning around 10am, and once in the afternoon, around 1pm, to capture bees across the spectrum of diurnal activity. On sampling days, we recorded all flowering species in bloom within the site, approximately a hectare in size, to provide an estimate of floral richness and seasonal bloom progression. We also used an infrared thermometer to record surface temperatures of three different external branches of each plant at noon on sampling days to test for effects of potentially warmer, darker plants.

### Specimens Processing & Data Management

All bees were labeled and pinned into field boxes each evening, then frozen for 48 hours to protect from insect infestations, and transported to Utah where they were identified to described species or unique morphospecies by experts at the USDA-ARS Pollinating Insect Research Unit (“Logan Bee Lab”). Bee identifications were completed using high quality ‘Leica’ dissecting microscopes, the appropriate taxonomic keys where available, and confirmed by comparison with the Logan Bee Lab's extensive reference collection of approximately 1.5 million specimens. Bees were assigned unique matrix code numbers that were included with standard insect label data printed on labels affixed to each specimen pin. The unique identifier and specimen field data were captured in a mySQL relational database, which was then managed and queried for statistical analyses using Microsoft Access front end software.

### Statistical Analyses

We employed a generalized linear mixed effects model with a negative binomial distribution to assess differences in bee visitation rates among the different plant treatments, which were modeled as fixed effects. We also used fixed effects to control for linear changes in visitation rates over the course of the day and differences in average visitation rates among the three sites. Variation in average visitation rates among the 63 plants and among the 9 sampling dates were each accounted for as a random effect. Because we intentionally collected on warm, sunny, calm days, the variation in environmental variables was minimal and their inclusion in the statistical model did not change treatment significance. They were therefore omitted from the final model for clarity.

Differences among treatments were estimated by comparing the number of bees that would be expected to visit a given plant in a given five-minute observation window under different conditions according to the negative binomial model. Ninety-five percent confidence intervals for the effect of each treatment versus the control were calculated using the model's variance-covariance matrix (Lawless 1987). To assess the possible tendency for bees to cluster on individual plants at a given point in time beyond what would be expected by treatment effects, we used a likelihood ratio test to compare one version of this model with an overdispersed error distribution (the negative binomial) to a version without overdispersion (the Poisson) (Coxe et al. 2009).

To compare the branch temperatures between blackened and not blackened branches, we built a linear mixed effects model with branch color (blackened or not) and day of year as fixed effects, and the plant within the site as a random effect (Fig. A1). All analyses were performed in the R programming language using the *lme4* package (Bates et al. 2015, p. 4; R Core Team 2015). All data and code used are freely available by contacting the corresponding author.

## Results

### Bee Collection

Despite a lack of floral cues, our 378 plant samples yielded 308 bees from forty-four different species across nine genera and five of the six North American bee families (Table 1). Approximately three-quarters of this bee abundance and diversity came from the two sprayed sugar treatments (N=220, Species=38). Shrubs with naturally-occurring mold, and hence honeydew sugars, attracted more bees and species (N=41, Spp.=15), than any of the three treatments not anticipated to be attractive to bees (*Control* N=11, Spp.=4; *Insecticide* N=17, Spp.=11; *Natural Mold + Insecticide* N=12, Spp.=8), or the treatment designed as a visual mimic of the dark color of mold (*Paint* N=7; Spp.=4).

**Table 1:**
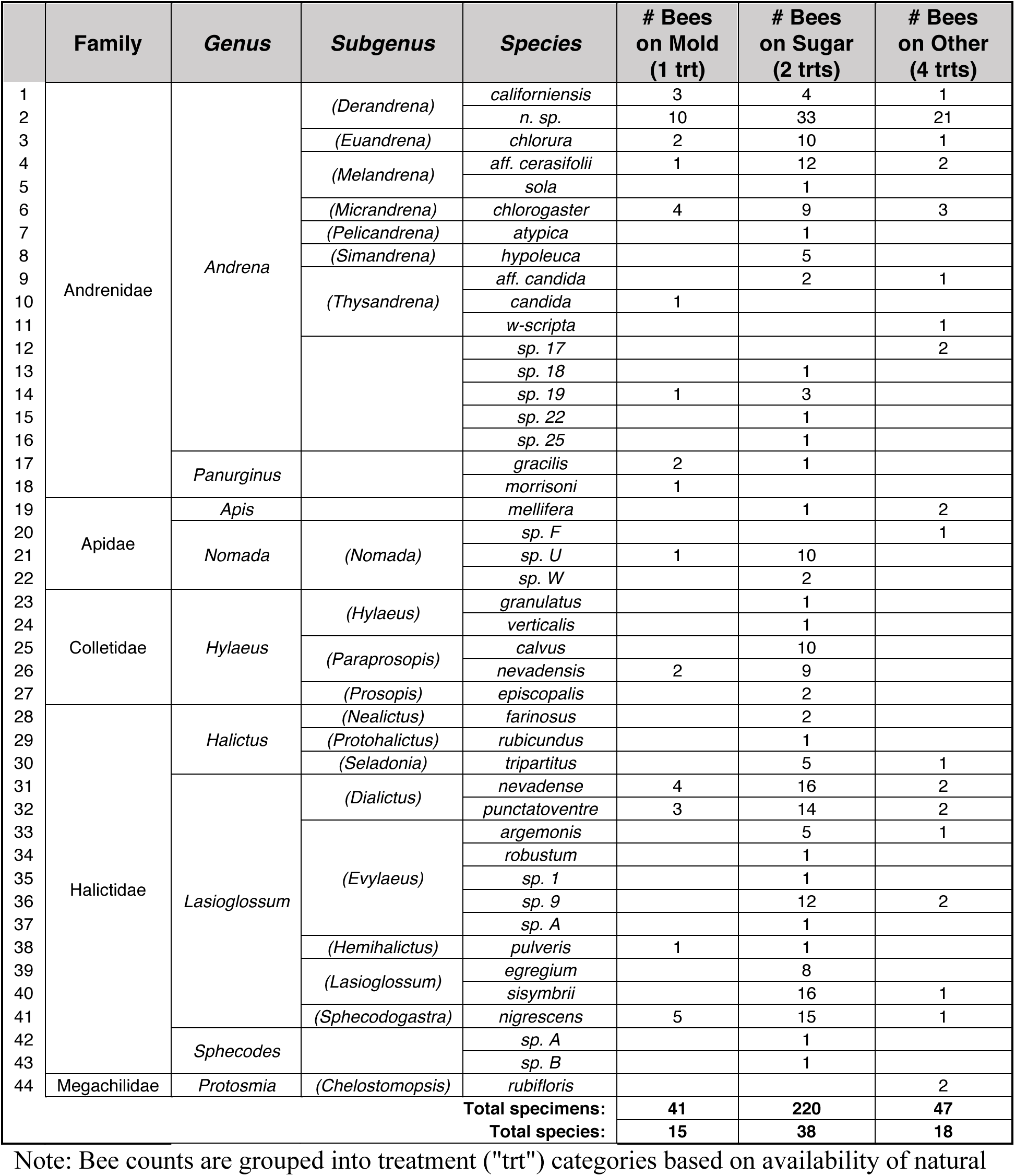
Faunal list and count of bees collected on treated non-flowering *A. fasciculatum*

Note: Bee counts are grouped into treatment ("trt") categories based on availability of natural sugars from honeydew (*Natural Mold)*, sprayed honeydew-mimic sugars (*Sugar, Sugar + Paint)*, or no known sugars accessible (*Control, Insecticide, Mold + Insecticide, Paint*).

### Seasonal and Spatial Context

Floral richness increased linearly across the season and across 1-hectare experimental sites as expected, from zero to thirteen species recorded in bloom during sampling, confirming that sampling captured bee activity during the relatively nectar-depauperate period leading up to peak bloom (which typically occurs at much higher richness and abundance at Pinnacles than was observed during the study). Likewise, total bee specimens collected increased over the nine-week duration of the study at all three sites, from the first sampling round (N=85) to approach peak bee activity during the third and final sampling round (N=146). Bee abundance differed somewhat between sites, with bee activity at sites C (N=125) and B (N=115) consistently higher than bee activity at site A (N=68). None of these temporal or site variables, however, nor any of the environmental variables recorded (e.g. cloud cover, ambient temperature, wind speed, humidity) influenced the modeled significance of treatment results.

### Treatment Significance

Our model results confirmed our original observation that native bee visitation to pre-bloom *Adenostoma fascicultum* is significantly elevated on plants with sooty mold, despite this resource lacking any floral cue (*p*=0.02). Model results also revealed our unadvertised honeydew-mimic solution to be significantly more attractive to bees than mold (*p*<0.001; Fig. 3), identifying simple sugars as the resource of interest in these nectar-poor landscapes. Furthermore, though there was no base effect of the *Insecticide* treatment on bee visitation rate compared to the *Control* (*p*=0.38, Fig. 4), a significant interaction between the *Mold* and *Insecticide* treatments reflects lower bee counts on moldy plants on which insecticide was applied to stop the production of honeydew (*p*=0.04; Fig. 4), indicating that active sugar production by live scale insects was a greater attraction to bees than residual sugars on branches or any visual or olfactory cue from scale insect carapaces. Bees were also not using the dark color of mold as a cue to locate honeydew, as evidenced by the lack of significant bee visitation effects of the *Paint* treatment (*p*=0.44; Fig. 3) or any interaction between the *Sugar* and *Paint* treatments (*p*=0.91; Fig. 4). Finally, since branch infrared thermometer readings did not differ between treatments (*p*=0.55; Fig. A1), observed bee behaviors can also not be explained by a response to thermal cues. From this experiment, we conclude that a highly diverse array of native, mostly non-social bees are visiting pre-bloom *Adenostoma fasciculatum* shrubs for sugars gleaned from honeydew, and are able to do so using foraging strategies outside the current framework centered around floral displays.

**Figure 3:**
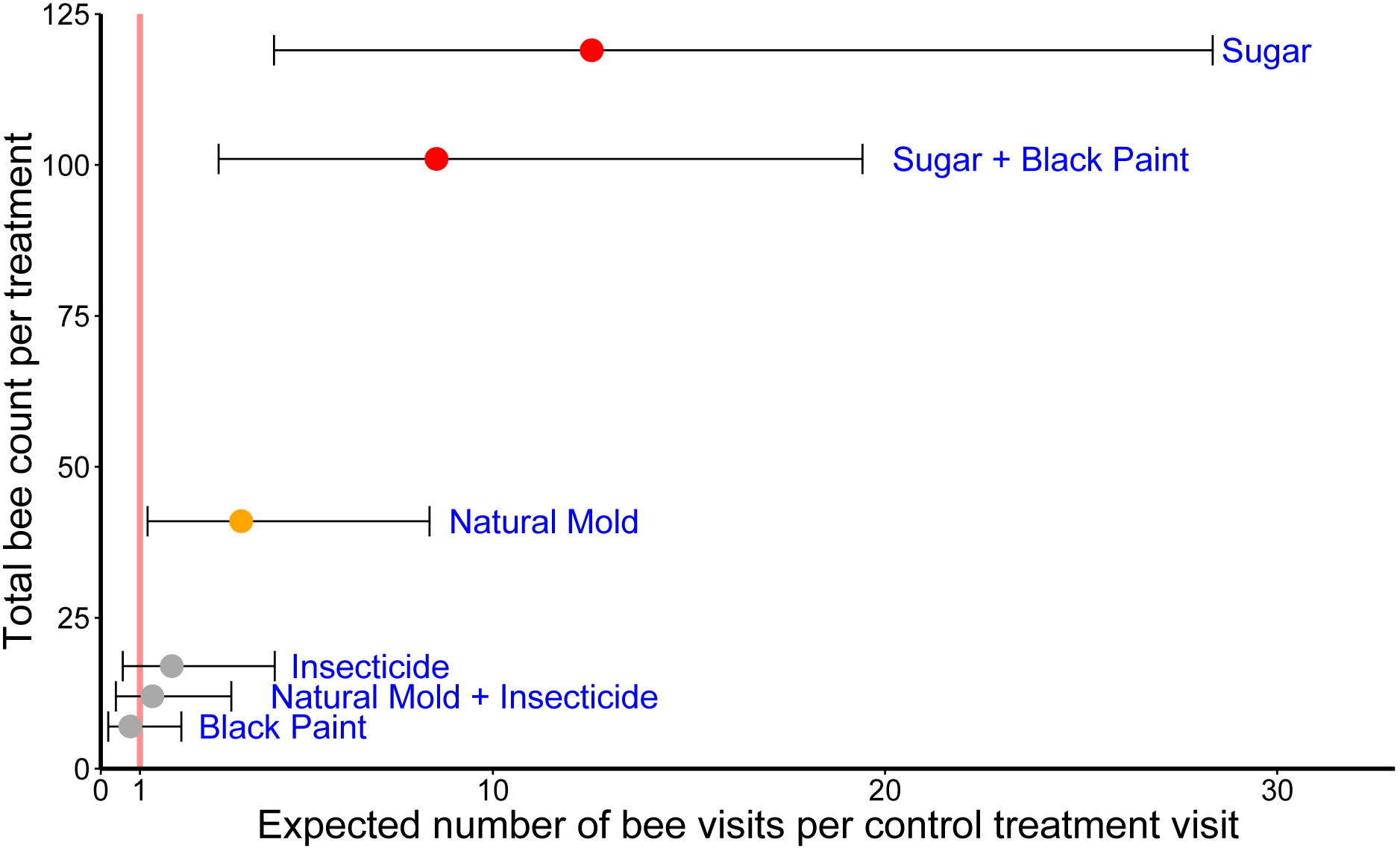
Effect sizes of bee visitation with 95% confidence intervals (x-axis) and total bee counts (y-axis) for each of six experimental plant treatments (labeled in blue) compared to the control treatment value (red vertical line). According to our generalized linear mixed effects model, the presence of *Natural Mold* increased the expected bee visitation rate by an estimated factor of 4 over the *Control* (*p* = 0.02, orange dot). Bee visitation to both sprayed *Sugar* treatments, even in the absence of any obvious cues, was significantly higher than that to *Natural Mold* (*p* < 0.001), and higher by an estimated factor of 13 over the *Control* (*p* < 0.0001, red dots). The three treatments without natural or sprayed sugars did not differ in bee visitation from the *Control* (*p* > 0.05, grey dots, 95% CIs overlapping the red line representing the *Control* treatment value). Observed counts on the y-axis are based on 54 observations of each treatment, divided evenly among the three sites.

**Figure 4:**
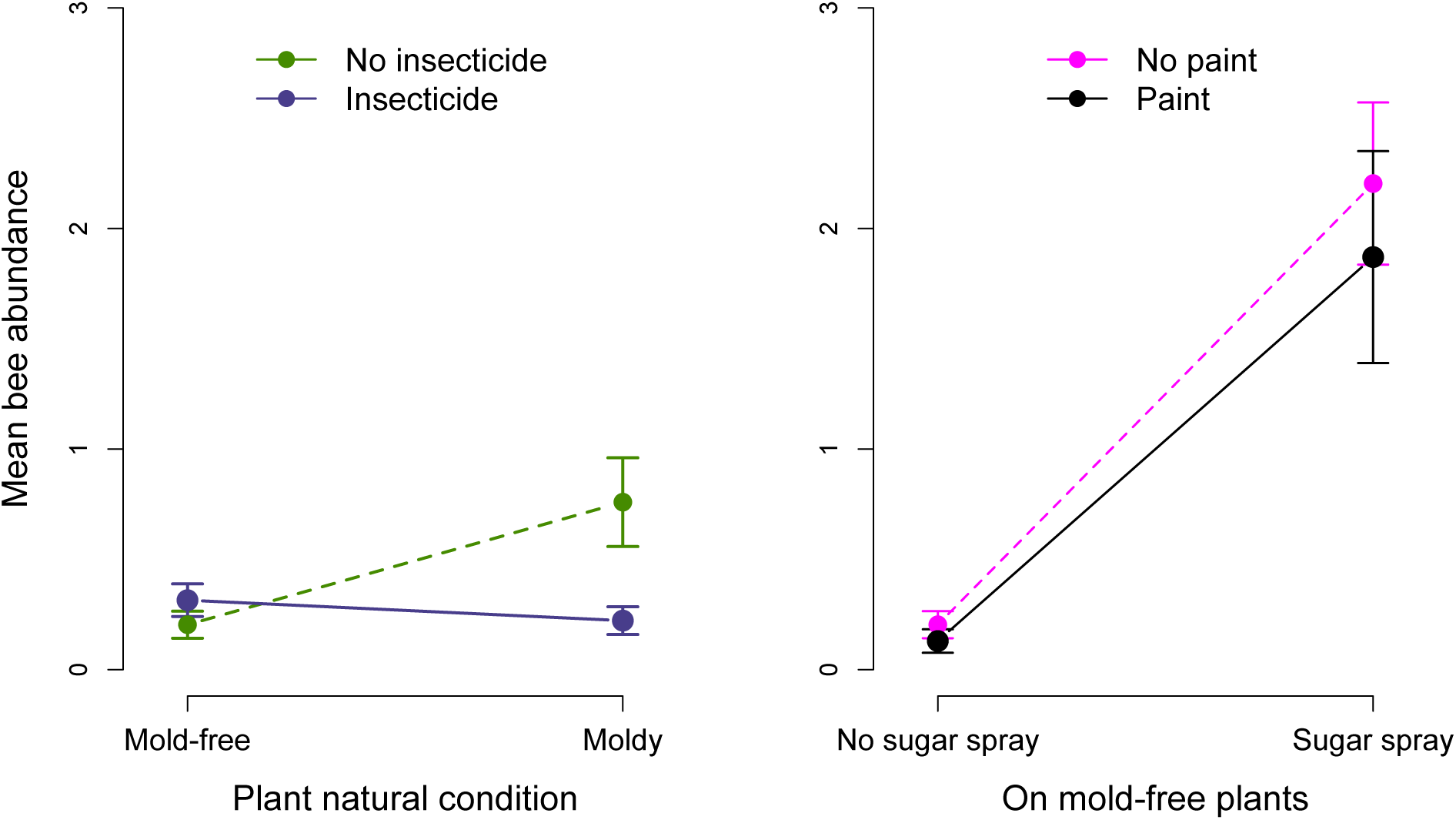
Two-way interactions of mean sample bee abundance between key treatments. A negative-binomial generalized linear mixed effects model with random intercepts for individual plants and for individual sampling dates found a significant interaction between the *Natural Mold* and *Insecticide* treatments (left, *p*=0.04), but not between the *Sugar* and *Black Paint* treatments (right, *p*=0.91). Error bars represent +/− one s.e.m. Number of samples and total bee counts per treatment are as reported in Figure 3.

## Discussion

Our study is the first to document the use of honeydew as a sugar resource across a diverse community of native bees as well as the first to bring to light widespread sugar foraging behaviors seemingly divorced from floral cues. Overall, we recorded forty-four bee species in nine genera and five of the six North American bee families exhibiting foraging patterns largely outside the general understanding that bee search images are behaviorally and evolutionarily tied to elaborate floral displays. Thirty-eight species of these native, mostly solitary bees were accessing our honeydew-mimic sugars sprayed on inconspicuous, non-flowering shrubs that offered no other reward, fifteen species of bees visited pre-bloom plants for natural honeydew absent any floral signal, and eighteen bee species displayed non-floral-centric foraging behaviors on other plant treatments (Table 1).

These results raise the question: how are so many species of bees rapidly locating sugar sprayed on a stick? Our current understanding of sensory abilities in wild bees does not explain this phenomenon. While it remains possible that bees are independently finding these plants via some unknown sensory cue from the sugar, our data distribution (Fig. 2A) includes several very high values on *Sugar* treatment plants (up to 22 bees in five minutes) that are more compatible with non-independent arrivals (Lawless 1987). Indeed, our data fit a negative binomial model that includes clustering on individual plants in specific 5-minute periods much better than the Poisson model that assumes independent arrivals (χ^2^=30, df=1, p<0.0001) (Coxe et al. 2009).

One way to explain this non-independence might be a strategy by which solitary bees are locating nontraditional sugars using social cues from other bee foragers, in combination with stochastic exploration of resources outside the floral realm. Social foraging mechanisms, such as the honeybee waggle dance (intraspecific) or the relationship between scavengers and primary predators (interspecific), are well-known for many animals, especially when resources are variable across space and time (Stahler et al. 2002; Deygout et al. 2010; Frisch 2014).

Interspecific foraging dynamics among bees, however, are not understood. For solitary bees that must provision a nest for offspring without the help of nest mates, cueing off the activity of heterospecifics in their community to opportunistically harvest unusual sugar resources could help optimize energetically-expensive foraging flights, especially in early-season or degraded habitats lacking sufficient bloom. Clearly, more research into these patterns and the ability of bees to locate non-floral sugars is required.

Regardless of the mechanism by which bees are able to find honeydew secretions, this behavior displayed by so many different wild bee species may have important implications for how bees will respond to a changing world with increasingly unpredictable conditions.

Mediterranean habitats, where bees are most diverse, have been identified as particularly vulnerable to climate change, exotic species invasions, and urbanization (Michener 2007; Klausmeyer and Shaw 2009). Warming temperatures have been found to cause shifts in the emergence time of solitary bees in relation to their preferred host plants, resulting in a temporal decoupling of plants from their pollinators (Inouye 2008; Bartomeus et al. 2011; Forrest and Thomson 2011; Robbirt et al. 2014). Ongoing habitat loss, fragmentation, and degradation are also threats to wild bee species (Steffan-Dewenter et al. 2002; Fahrig 2003; Cane et al. 2006), many of which are only active for one month out of the year and rely on their preferred pollens being available during that time (Linsley 1958). For bees that emerge during the early season into a habitat of unexpectedly poor floral resources, the ability to subsist on alternate sugar sources that would extend longevity until nectar and pollens can be located could be critical to survival and production of offspring.

In conclusion, the occurrence of over forty different species of native bees on an unadvertised, non-floral sugar resource suggests widespread, previously undocumented plasticity in bee foraging behaviors and diet breadth that may become increasingly relevant to bee conservation with continued disruptions in floral bloom. This discovery represents not only a novel behavioral phenomenon and notable departure from the historical focus on bee use of visual, olfactory, and floral cues, but may also have implications related to both the resilience of bee communities to temporary habitat perturbations and the social complexity of their foraging dynamics. Our finding that diverse solitary bees use nontraditional resources and foraging strategies during times of low bloom suggests that bee use of honeydew may be only one example of adaptive bee foraging strategies that have yet to be described. Future research on native bee foraging behaviors may benefit from considering the effect of stochastic and socially-mediated foraging behaviors, and investigating the use of non-floral, unadvertised resources.

## Acknowledgments

The authors are grateful to Pinnacles National Park for funding this project through the Great Basin Cooperative Ecosystem Studies Unit (Task Agreement number P10AC00577). JMM is currently supported by a University of Florida Biodiversity Institute Graduate Fellowship, and previously by the Utah State University Ecology Center. DJH is supported by Gordon and Betty Moore Foundation's Data-Driven Discovery Initiative Grant GBMF4563 to E.P. White. We owe thanks to Paul Johnson for on-site guidance, photography, and field errands, Edward W. Evans for honeydew discussions and feedback on early drafts, J. Therese Lamperty for diligent fieldwork, H. Ikerd and S. Burrows for meticulous lab work, G. Yenni, A. Kleinhesselink and S. Durham for statistical advice, and the Weecology lab group for manuscript feedback. JMM would also like to thank Dr. Yael Mandelik for hosting her for a related field project at the Hebrew University of Jerusalem in Rehovot, Israel, which was funded by a ‘Rahamimoff Travel Grant for Young Scientists’ from the US-Israel Binational Science Foundation.

## Author Contributions

JMM initiated and designed the project, executed the field work and analyses, and wrote the paper. TLG oversaw bee identifications and advised on protocol. DJH improved the statistical analyses and assessed statistical significance. SKME guided analyses, concept, and writing. All authors discussed results and commented on the manuscript.

## Additional Information

Upon article publication, data for this project will be available at http://www.datadryad.org. Code and data are currently available online at https://github.com/beecycles/Bees-without-flowers-project2016 and are citable using doi.org/10.5281/zenodo.162054. Correspondence and requests for materials should be addressed to JMM.

## Expanded Online Materials

### Online Appendix A: Additional Methodological and Data Details

**Table A1:** Summary of plant types and treatment sprays used in seven experimental treatments.

| Natural Plant Condition | Treatment Number | Treatment Description | Treatment Spray applied to each plant |
| --- | --- | --- | --- |
| Moldy | Moldy (i) | Natural Mold | 5ml distilled water |
|  | Moldy (ii) | Natural Mold + Insecticide | 5ml Orange Guard® short-residual insecticide |
| Mold-free | Mold-free (i) | Control | 5ml distilled water |
|  | Mold-free (ii) | Insecticide | 5ml Orange Guard® short-residual insecticide |
|  | Mold-free (iii) | Black Paint | 5ml non-toxic black paint |
|  | Mold-free (iv) | Sugar | 5ml 20% chemical-grade Fructose:Sucrose |
|  | Mold-free (v) | Sugar + Black Paint | 5ml 20% chemical-grade Fructose:Sucrose + 5ml non-toxic black paint |

Note: Each treatment was replicated with three plants at three different sites, for a total of nine plants per treatment and sixty-three plants total. Each site was sampled in the morning and afternoon on three separate occasions.

**Figure A1:**
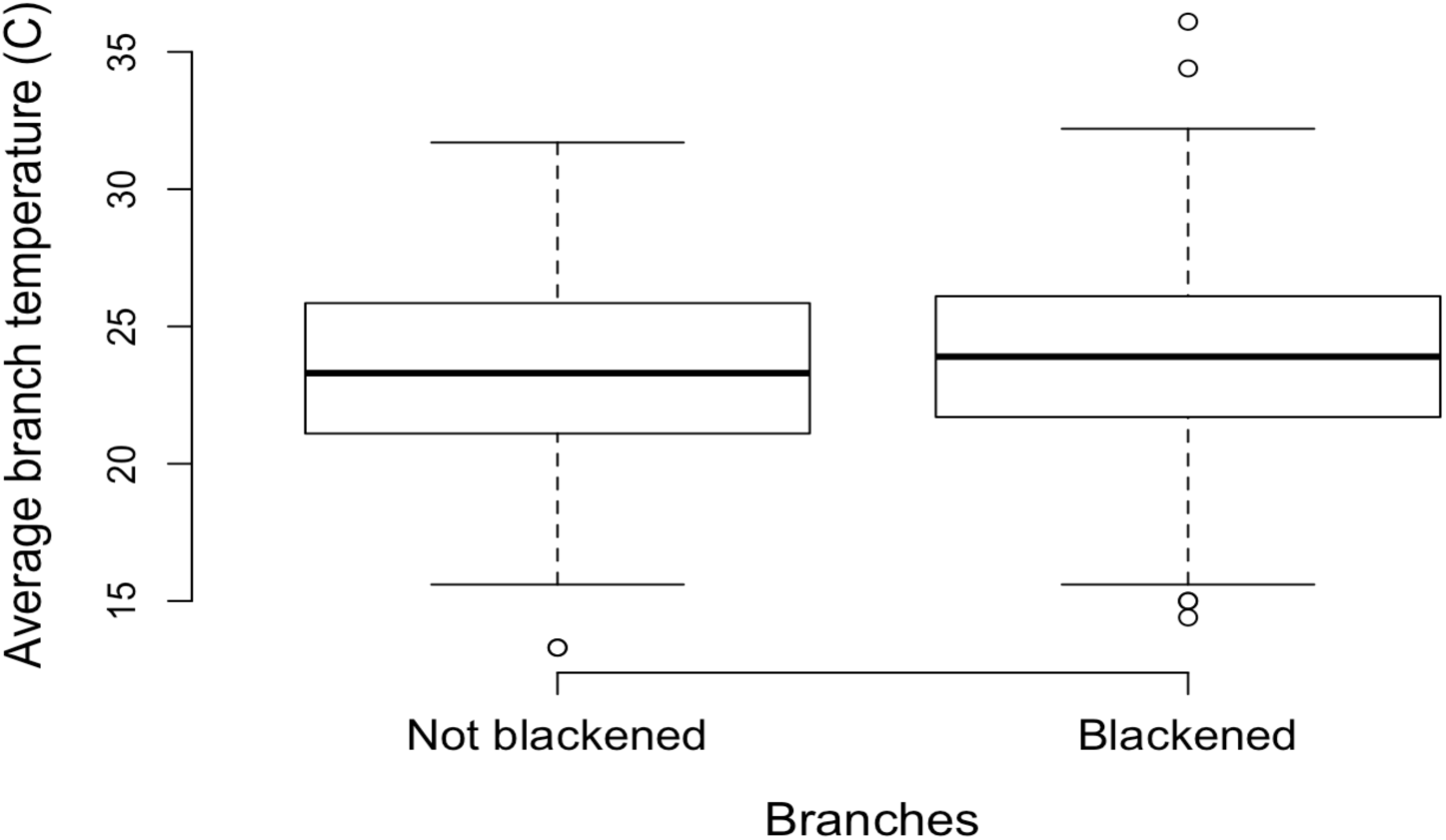
Surface temperatures (C°) for “blackened” and “not blackened” branches of treatment plants, measured at noon with an infrared thermometer on each of six sampling days. A linear mixed effects model that controlled for day of year as a fixed effect and plant within site as a random effect found no difference in surface temperatures between blackened (N=324) and not blackened (N=243) branches (*p*=0.55).

**Figure A2:**
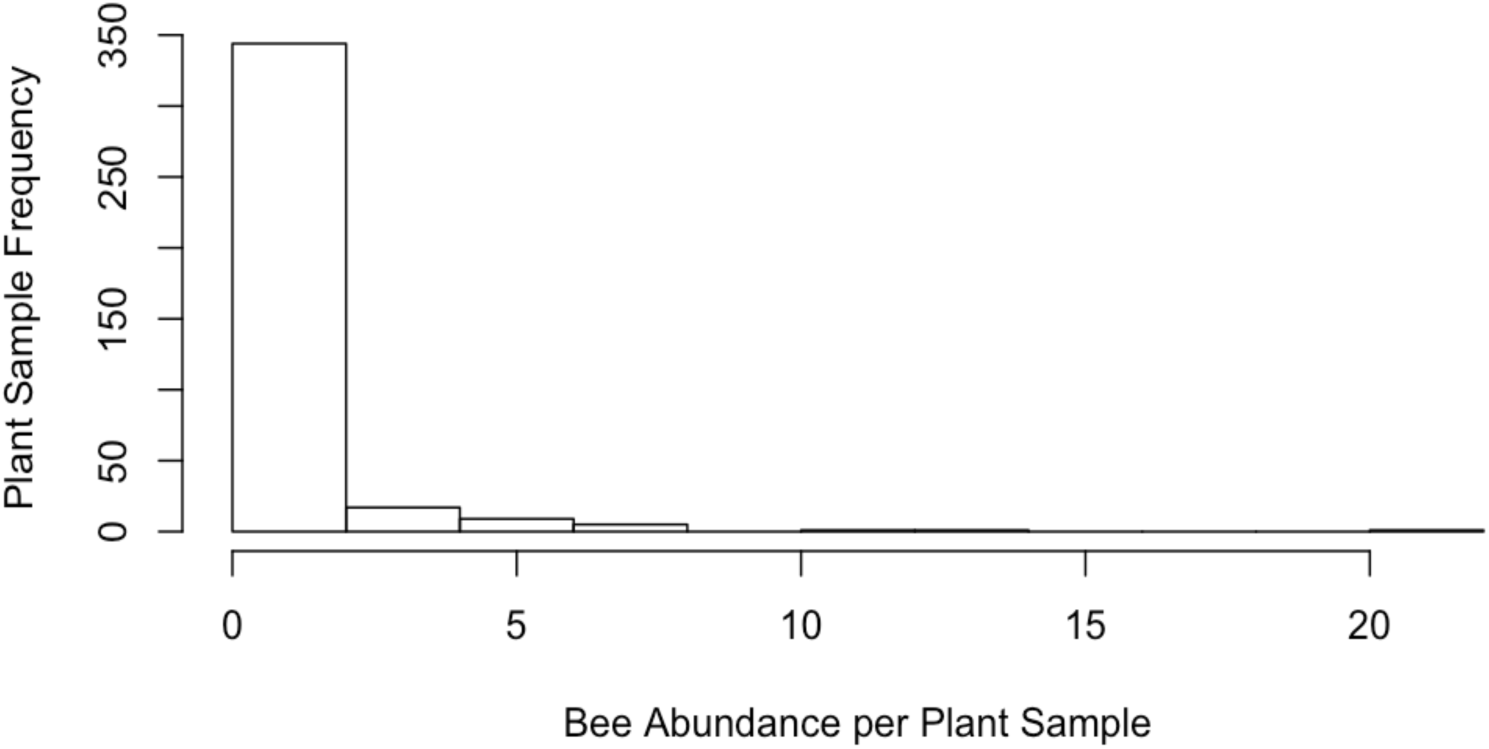
Negative binomial distribution of bee count response variable (N=308, from 378 samples at treatment plant). Bee visitors to experimental plants were collected with an aerial net during randomly-ordered five-minute sampling periods at each plant. Many periods of zero bees collected and a few events where up to 22 bees were collected on an experimental plant produce this data distribution, which limited statistical analyses to the tests described in methods. Since biological count data are typically Poisson-distributed and do not include the high values we see on the right side of this histogram from bees visiting *Sugar* treatment plants (N = 22, 14, 11, 8, 8, 8), we point to these select high values as possible evidence of feedback from a social foraging mechanism, and encourage further research in this area (Lawless 1987).

## Literature Cited

Bartomeus, I., J. S. Ascher, D. Wagner, B. N. Danforth, S. Colla, S. Kornbluth, and R. Winfree. 2011. Climate-associated phenological advances in bee pollinators and bee-pollinated plants. Proceedings of the National Academy of Sciences 108:20645–20649.

Bates, D., M. Maechler, B. Bolker, and S. Walker. 2015. Fitting Linear Mixed-Effects Models Using lme4. Journal of Statistical Software 67:1–48.

Batra, S. W. T. 1993. Opportunistic Bumble Bees Congregate to Feed at Rare, Distant Alpine Honeydew Bonanzas. Journal of the Kansas Entomological Society 66:125–127.

Bishop, J. A. 1994. Bumble Bees (Bombus hypnorum) Collect Aphid Honeydew on Stone Pine (Pinus pumila) in the Russian Far East. Journal of the Kansas Entomological Society 67:220–222

Bukovac, Z., A. Dorin, V. Finke, M. Shrestha, J. Garcia, A. Avarguès-Weber, M. Burd, et al. 2016. Assessing the ecological significance of bee visual detection and colour discrimination on the evolution of flower colours. Evolutionary Ecology 1–20.

Cane, J. H., R. L. Minckley, L. J. Kervin, T. H. Roulston, and N. M. Williams. 2006. Complex Responses Within A Desert Bee Guild (Hymenoptera: Apiformes) To Urban Habitat Fragmentation. Ecological Applications 16:632–644.

Chittka, L., N. M. Williams, H. Rasmussen, and J. D. Thomson. 1999. Navigation without vision: bumblebee orientation in complete darkness. Proceedings of the Royal Society of London. Series B: Biological Sciences 266:45–50.

Clarke, D., H. Whitney, G. Sutton, and D. Robert. 2013. Detection and Learning of Floral Electric Fields by Bumblebees. Science 340:66–69.

Coxe, S., S. G. West, and L. S. Aiken. 2009. The Analysis of Count Data: A Gentle Introduction to Poisson Regression and Its Alternatives. Journal of Personality Assessment 91:121–136.

Crane, E., and P. Walker. 1985. Important Honeydew Sources and their Honeys. Bee World 66:105–112.

de Jager, M. L., L. L. Dreyer, and A. G. Ellis. 2011. Do pollinators influence the assembly of flower colours within plant communities? OECOLOGIA 166:543–553.

Deygout, C., A. Gault, O. Duriez, F. Sarrazin, and C. Bessa-Gomes. 2010. Impact of food predictability on social facilitation by foraging scavengers. Behavioral Ecology 21:1131–1139.

Dyer, A. G., H. M. Whitney, S. E. J. Arnold, B. J. Glover, and L. Chittka. 2006. Bees associate warmth with floral colour. Nature 442:525–525.

Fahrig, L. 2003. Effects of Habitat Fragmentation on Biodiversity. Annual Review of Ecology, Evolution, and Systematics 34:487–515.

Forrest, J. R. K., and J. D. Thomson. 2011. An examination of synchrony between insect emergence and flowering in Rocky Mountain meadows. Ecological Monographs 81:469–491.

Friel, E. N., R. S. T. Linforth, and A. J. Taylor. 2000. An empirical model to predict the headspace concentration of volatile compounds above solutions containing sucrose. Food Chemistry 71:309–317.

Frisch, K. von. 2014. Bees: Their Vision, Chemical Senses, and Language. Cornell University Press.

Herrera, C. M. 1995. Floral Biology, Microclimate, and Pollination by Ectothermic Bees in an Early-Blooming Herb. Ecology 76:218–228.

Inouye, D. W. 2008. Effects of climate change on phenology, frost damage, and floral abundance of montane wildflowers. Ecology 89:353–362.

Klausmeyer, K. R., and M. R. Shaw. 2009. Climate Change, Habitat Loss, Protected Areas and the Climate Adaptation Potential of Species in Mediterranean Ecosystems Worldwide. PLoS ONE 4:e6392.

Koch, H., C. Corcoran, and M. Jonker. 2011. Honeydew Collecting in Malagasy Stingless Bees (Hymenoptera: Apidae: Meliponini) and Observations on Competition with Invasive Ants. African Entomology 19:36–41.

Konrad, R., F. L. Wäckers, J. Romeis, and D. Babendreier. 2009. Honeydew feeding in the solitary bee Osmia bicornis as affected by aphid species and nectar availability. Journal of insect physiology 55:1158–1166.

Lawless, J. F. 1987. Negative binomial and mixed Poisson regression. Canadian Journal of Statistics 15:209–225.

Linsley, E. G. 1958. The Ecology of Solitary Bees. Hilgardia 27:543–599.

Michener, C. D. 2007. The Bees of the World. Johns Hopkins University Press, Baltimore.

Ohashi, K., and T. Yahara. 2001. Behavioural responses of pollinators to variation in floral display size and their influences on the evolution of floral traits. Pages 274–296 *in*Cognitive Ecology of Pollination. Cambridge University Press.

Ollerton, J., R. Winfree, and S. Tarrant. 2011. How many flowering plants are pollinated by animals? Oikos 120:321–326.

Orbán, L. L., and C. M. S. Plowright. 2014. Getting to the start line: how bumblebees and honeybees are visually guided towards their first floral contact. Insectes Sociaux 61:325–336.

Potts, S. G., J. C. Biesmeijer, C. Kremen, P. Neumann, O. Schweiger, and W. E. Kunin. 2010. Global pollinator declines: trends, impacts and drivers. Trends in Ecology & Evolution 25:345–353.

R Core Team. 2015. R: A language and environment for statistical computing. https://www.Rproject.org. Vienna, Austria.

Regal, P. J. 1977. Ecology and Evolution of Flowering Plant Dominance. Science 196:622–629.

Robbirt, K. M., D. L. Roberts, M. J. Hutchings, and A. J. Davy. 2014. Potential Disruption of Pollination in a Sexually Deceptive Orchid by Climatic Change. Current Biology 24:2845–2849.

Santas, L. 1983. Insects producing honeydew exploited by bees in Greece. Apidologie 14.2:93–103.

Stahler, D., B. Heinrich, and D. Smith. 2002. Common ravens, Corvus corax, preferentially associate with grey wolves, Canis lupus, as a foraging strategy in winter. Animal Behaviour 64:283–290.

Steffan-Dewenter, I., U. Münzenberg, C. Bürger, C. Thies, and T. Tscharntke. 2002. Scale-Dependent Effects of Landscape Context on Three Pollinator Guilds. Ecology 83:1421–1432.

Thorp, R. W., D. L. Briggs, J. R. Estes, and E. H. Erickson. 1975. Nectar Fluorescence under Ultraviolet Irradiation. Science 189:476–478.

Wäckers, F. L. P. C. J. van Rijn, and G. E. Heimpel. 2008. Honeydew as a food source for natural enemies: Making the best of a bad meal? Biological Control, Conservation Biological Control 45:176–184.

Whitney, H. M., A. Dyer, L. Chittka, S. A. Rands, and B. J. Glover. 2008. The interaction of temperature and sucrose concentration on foraging preferences in bumblebees. Naturwissenschaften 95:845–850.

Wright, G. A., and F. P. Schiestl. 2009. The evolution of floral scent: the influence of olfactory learning by insect pollinators on the honest signalling of floral rewards. Functional Ecology 23:841–851.

Zoebelein, G. 1957. Die Rolle des Waldhonigtaus im Nahrungshaushalt forstlich nützlicher Insekten. Forstwissenschaftliches Centralblatt 76:24–34.

